## Supplementary data 1 for "Selective attention and sensitivity to auditory disturbances in a virtually-real Classroom: Comparison of adults with and without AD(H)D"

**Supplementary Materials**

Univariate vs. Multivariate TRF estimation

The soundscape in the Events condition contained a combination of the teacher’s speech and the occasional sound-events played in the background. In the main manuscript we report the TRF estimated to capture the response to the teacher’s speech using a multivariate model, that takes into account both the acoustic envelope of the speech as well as the sound-events. Here we compared TRF estimation using this multivariate encoding model vs. a univariate encoding model comprised only the speech-envelope as a single regressor.

Figure S1A shows that the TRFs estimated in response to the speech were qualitatively similar using both approaches, with a negative peak around 100ms followed by a positive peak at 150ms, similar to the pattern observed in previous studies (Lalor and Foxe, 2010; Zion Golumbic et al., 2013; Ding and Simon, 2014; Kaufman and Zion‐Golumbic, 2023). These TRFs were also qualitatively similar to the TRF estimated in the Quiet condition, which did not contain additional event-sound, lending additional credibility to the approach. In the multivariate model, a TRF capturing the response to sound-events was also estimated (Figure S1B), which also shows two prominent peaks - negative at 100ms and positive at 200 - a pattern comparable to the ERPs derived for these stimuli through simple averaging, as expected for transient stimuli (given the linear nature of the model). When comparing the predictive power of the univariate and multivariate models, both models yielded significant predictive power relative to null-permutations, however the multivariate model was able to explain a larger degree of the variance in the neural response [t(48) = 10.354, p < 0.0001, Cohen’s d = 1.268, BF10 = +100 (Extreme evidence for H1); Figure S1C].

These results indicate that including a regressor that captures the time-course of the sound-events in addition to a regressor capturing the speech stimulus allowed the model to more faithfully capture the neural representation of the entire VR classroom soundscape (Crosse et al., 2021). Therefore, this approach is preferable when analyzing neural response to stimuli embedded in realistically noisy contexts. That said, the spatio-temporal TRF itself can be reliably extracted for the speech stimulus using both a univariate and a multivariate model.

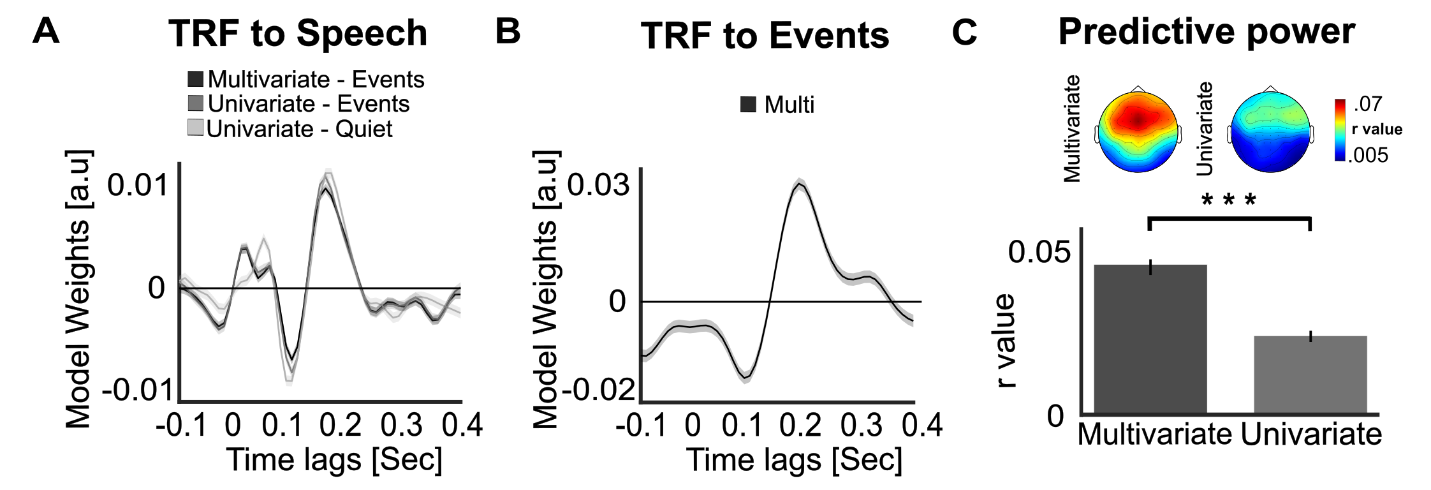

**Figure S1. Comparison between Univariate and Multivariate TRF model. A.** Grand-average TRFs (across both groups) reflecting the neural tracking response of the teacher’s speech, estimated for the Events condition using both a univariate and multivariate model, as well as for the Quiet condition (univariate model). Shading represent the standard error of the mean (SEM) **B.** Grand-average TRF estimated for the regressor representing the audio of the event-sounds, in the multivariate model of the Events condition. Shaded areas represent the SEM. This analysis is akin to calculating the event-related potential (ERP) to event-sounds. **C.** Topographical distribution (top) and bar graph (bottom) depicting the predictive power values (Pearson’s r) in the Events condition for the univariate and multivariate encoding models.
